# How the dominant reading direction changes parafoveal processing: A combined EEG/eye-tracking study

**DOI:** 10.1101/2023.01.30.526189

**Authors:** Xin Huang, Hezul Tin-Yan Ng, Chien Ho Lin, Ming Yan, Olaf Dimigen, Werner Sommer, Urs Maurer

## Abstract

Reading directions vary across writing systems. Through long-term experience readers adjust their visual systems to the dominant reading direction in their writing systems. However, little is known about the neural correlates underlying these adjustments because different writing systems do not just differ in reading direction, but also regarding visual and linguistic properties. Here, we took advantage that Chinese is read to different degrees in left-right or top-down directions in different regions. We investigated visual word processing in participants from Taiwan (both top-down and left-right directions) and from mainland China (only left-right direction). Combined EEG/eye tracking was used together with a saccade-contingent parafoveal preview manipulation to investigate neural correlates, while participants read 5-word lists. Fixation-related potentials (FRPs) showed a reduced late N1 effect (preview positivity), but this effect was modulated by the prior experience with a specific reading direction. Results replicate previous findings that valid previews facilitate visual word processing, as indicated by reduced FRP activation. Critically, the results indicate that this facilitation effect depends on experience with a given reading direction, suggesting a specific mechanism how cultural experience shapes the way people process visual information.

## 1. Introduction

A neurocognitive framework that the brain and mind are shaped by sociocultural experience has gained much attention (Han and Northoff, 2008). One aspect of sociocultural experience, the writing system and the reading direction are considered to be important in shaping the neurocognitive networks (Kazandjian and Chokron, 2008). Cultures have developed different writing systems that vary in terms of their reading direction. For example, Roman alphabetic languages are written and read from left to right whereas Hebrew and Arabic Abjad are read from right to left. The Ancient Greek “boustrophedon” writing style alternated between left-right and right-left directions, like the ox turns when plowing a field. Although modern Chinese is most frequently written from left to right, it was traditionally written in top-down direction, and many Chinese readers are still regularly exposed to the top-down direction when reading novels or classical texts. Thus, learning to read Chinese also entails some familiarity with reading in top-down direction. Interestingly, how the visual system deals with such differences in reading direction, and whether it undergoes general adaptations during learning to read, is largely unknown.

Reading experience can influence the size of the perceptual span, that is, the area from which readers pick up information during a fixation (McConkie and Rayner, 1975; for review, see Rayner et al., 2009). An asymmetric perceptual span, being wider in the direction of reading than against, has been found in many writing systems. For example, skilled readers of English who read from left to right obtain useful information from an area extending 14–15 letter spaces to the right of fixation but only 3–4 letter spaces to the left (e.g., McConkie and Rayner, 1976; Rayner et al., 1980; Underwood and McConkie, 1985). Also in Japanese, where texts are written either in a horizontal or vertical direction, Osaka (1993) found that the perceptual span was asymmetric towards the direction of reading, depending on whether it was vertical or horizontal. When reading Chinese from left to right, the perceptual span extends one character to the left of fixation but two to four characters to the right (Inhoff and Liu, 1997; Yan et al., 2015). Therefore, it is established that readers’ perceptual span shows an asymmetrical pattern, extending further toward than against the direction of reading.

Two explanations have been suggested to account for this asymmetric perceptual span in reading. One is hemispheric specialization, which assumes that if the perceptual span is determined primarily by the hemispheric projections, then languages which are written from left-to-right should both produce a rightward asymmetry, as dominance for language is typically left-hemispheric (e.g., Almabruk et al., 2011; Ibrahim and Eviatar, 2012, 2009). An alternative explanation is reading experience, also known as the scanning effect, which assumes that a reader’s perceptual span is in line with the direction in which texts are read because of the experience with this typical reading direction. However, empirical studies have only supported reading experience accounts but provided no evidence for the hemispheric dominance account. Pollatsek et al. (1981) tested the extent of the perceptual span in native Israeli Hebrew/English bilinguals who read sentences while a gaze-contingent moving window extended either up to 14 characters to the left or to the right of fixation. Results showed that reading performance for Hebrew was superior when the asymmetric window was larger to the left, whereas performance for English was superior when the window was larger to the right. In addition, Jordan et al. (2013) provided evidence that the central perceptual span (an area extending 2.5 degrees to either side of fixation) was skewed according to the overall reading direction. In this study, skilled Arabic readers who were bilingual in Arabic and English read both Arabic and English sentences. In a symmetric window condition, the moving window of normal text extended 0.5 degrees to the left and right of fixation. In an asymmetric condition, the window was increased to 1.5 degrees or 2.5 degrees to either the left or the right. When reading English, performance across window conditions was superior when the window extended rightward. Conversely, when reading Arabic, performance was superior when the window extended leftward. Similar results were replicated for Urdu-English bilingual readers (Paterson et al., 2014) and in another left-running Arabic-based script. Thus, Zhou et al. (2021) determined the perceptual span in Uighur to cover 5 previous letters to the right of a fixation, and 12 upcoming letters to the left. Culturally acquired directional scanning habits even extend to non-text reading, namely, picture naming and recall (Padakannaya et al., 2002). Based on these results, it is believed that reading experience in a specific direction can modify the asymmetry of perceptual span.

Besides the asymmetric perceptual span, reading experience can also influence the preferred viewing locations (PVL) within words (Rayner, 1979). Yan et al. (2014) found that in Uighur scripts with a right-to-left reading direction, readers showed a rightward shifted PVL, meaning that more visual information about the fixated word is projected into the readers’ left visual field. In contrast, in scripts that are written from left to right (e.g., English), the PVL is shifted to the left (Deutsch and Rayner, 1999).

Whereas most studies that support the reading direction account used horizontal texts to investigate the asymmetry of the perceptual span, the horizontal-vertical contrast may be a better condition to test the two accounts summarized above. If readers are used to reading in a vertical direction, they should show a larger perceptual span in vertical texts than readers who are used to reading horizontal texts. When comparing the perceptual span in different writing directions, most previous studies used different scripts for different reading directions (except for studies on Japanese and Mongolian, Osaka and Oda, 1991; Su et al., 2020). Therefore, the observed differences in the asymmetry of perceptual span in different reading directions are confounded with differences in the script systems. In the present study we will avoid such confounds by using Chinese script, which allows to assess the asymmetry of perceptual span in different reading directions with the same stimuli and the same participants.

Historically, Chinese sentences were written vertically in columns going from top to bottom, with each new column starting to the left of the preceding one. Only rather recently, the horizontal alignment with a rightward reading direction was adopted. The People’s Republic of China has adopted horizontal alignment since 1956 along with the simplified Chinese orthographic reform, although vertical alignment is still occasionally used (e.g., in older Chinese books). While the horizontal alignment has been adopted in math and science texts, vertical alignment is still common in novels, newspapers, and magazines in other Chinese-speaking countries like Taiwan, Hong Kong, and Macau, where also traditional Chinese characters are employed. As a result, both horizontal and vertical reading directions are familiar and efficient to readers of traditional Chinese. In contrast, readers of simplified Chinese (as used in mainland China) are more familiar with horizontal alignments. This provides an opportunity to investigate the effects of experience with different reading directions in the same language.

Yan et al. (2019) conducted the first systematic analyses of eye movements during reading horizontal and vertical text among Taiwanese readers, who were familiar with both reading directions. These authors found that the participants read sentences equally efficiently, and that PVL distributions were highly similar in the two directions. In addition, there was a tradeoff between longer fixation durations in vertical than in horizontal reading but better fixations closer to the word center. This study implies that reading experience could differentially influence visual processing of Chinese characters in the horizontal and vertical directions of reading.

Previous eye-tracking studies on traditional Chinese in Taiwan showed similar findings as from simplified Chinese in mainland China, even though their reading and writing directions are different. For example, phonological (Tsai et al., 2004), morphological (Yen et al., 2008), and semantic information (Tsai et al., 2012) could be accessed in the parafoveal area during horizontal reading, which is consistent with simplified Chinese in mainland China (Liu et al., 2002; Yan et al., 2012, 2009). Even in vertical reading, semantic information was still accessible parafoveally (Pan et al., 2022). However, given that the studies were limited to Taiwanese participants, the direct comparison with oculomotor behavior in mainland Chinese is still lacking.

Although there is evidence from eye-tracking studies that cultural experience influences, how readers process information in different reading directions, the neural correlates of these processes are not yet known because of the limitations of typical neuroimaging methods. Functional magnetic resonance imaging (fMRI) has high spatial resolution, but its limited temporal resolution makes it typically unsuitable for explorations on individual target words. Conversely, event-related potentials (ERP) have high temporal resolution but the pervasive eye movement artifacts and other problems, such as the overlap between the ERP components elicited by successive fixations have hindered natural reading studies for a long time. However, recent developments of ocular detection and correction from eye-tracking information allow to deal with the eye movement artifacts in EEG data (Dimigen et al., 2011). The technique of EEG/eye-tracking co-registration has been developed and frequently used in unconstrained viewing situations, including reading, as it allows readers to move their eyes freely. By recording both eye movements and EEG, it is possible to obtain complementing information in terms of temporal and spatial domains, as eye-tracking can tell us, where observers fixate their gaze, while EEG registers, when and how the brain responds to the information. By time-locking the EEG to fixation onset, fixation-related potentials (FRPs) can capture perceptual and cognitive processes at the current fixation. This method has now also been frequently combined with eye-gaze contingent paradigms to study reading, including the boundary paradigm (Rayner, 1975). In this paradigm, an invisible boundary is embedded in the text. Prior to crossing the boundary, a parafoveal preview stimulus is shown instead of the actual target word. Only when the reader’s gaze crosses the boundary, the preview stimulus is replaced by the target word. By manipulating the relationship between the preview stimulus and the target word, it is possible to study the types of information readers extract from the parafovea. Modulation of the perceptual span can also be assessed in the boundary paradigm. A larger identity preview effect indicates that more parafoveal information has been acquired during previous fixations, thus implying a larger perceptual span. For instance, Inhoff et al. (1989) demonstrated that more parafoveal information was obtained from text when reading normal words than words were letter-transformed. Similarly, a reduction of the preview effect has been reported, when pre-target words were infrequent (Henderson and Ferreira, 1990). In Chinese reading, Yan et al. (2010) reported a larger preview effect from the second post-boundary word (i.e., word N + 2), when word N + 1 was more frequent. More recently, Yan and Sommer (2019) demonstrated that emotionally negative foveal words bind more attention than neutral and positive words leading to a reduced N+2 preview effect.

Studies combining EEG and eye-tracking found a reduced negativity (termed “preview positivity”) in FRPs following valid as compared to invalid previews in a time window between 200 and 280 ms after fixating the target word N + 1 (e.g., Dimigen et al., 2012; Kornrumpf et al., 2016), which was maximal over the occipito-temporal scalp. While this effect has often been referred to as “N1 effect”, its time window and scalp distribution are similar to the late N1 or N250 component observed in masked priming studies (e.g., Holcomb and Grainger, 2007). Therefore, the neural mechanism may be interpreted as a facilitatory effect of repetition suppression (Dimigen et al., 2012). The preview positivity is not only observed in word list reading (Dimigen et al., 2012; Niefind and Dimigen, 2016), but also in natural sentence reading (Degno et al., 2019a, 2019b; Dimigen and Ehinger, 2021).

In contrast to the late N1 component, the early parts of the “N1 effect” have been less frequently reported or investigated. The early N1 effect has been found in visual word processing with unrelated stimuli eliciting larger negativities compared to repeated stimuli (Niefind and Dimigen, 2016; Kornrumpf et al., 2016; Degno et al., 2019a) but was absent in Dimigen et al. (2012) and Li et al. (2015). Similar to the late N1 effect, the early N1 effect is largest activation in occipito-temporal regions of the scalp. Compared to the preview positivity, the early N1 effects are usually smaller and less robust, and also less consistent. Also, there appears to be a tendency that the early N1 preview effects are larger in Chinese (i.e., Li et al., 2022b, 2022a, 2015) than in alphabetic languages (i.e., Dimigen and Ehinger, 2021), possibly because of the higher visual complexity of Chinese and higher demands on visual processing (McBride-Chang et al., 2011; Zhao et al., 2014).

The present study co-registered eye movements and EEG in the boundary paradigm to investigate the neural correlates underlying the preview effects in two participant groups that differ with regard to their experience with different reading directions but essentially use the same script system. To this end, we recruited participants from mainland China, where Chinese is written from left to right, and participants from Taiwan, where Chinese is written in both top-down and left-to-right directions but more often top-down. Both groups were tested with the same materials in both vertical and horizontal directions. Importantly, only characters were used as materials that are identical in simplified and traditional Chinese script.

For both, the early and late N1 components, we expected reduced (more positive) amplitudes after identical previews as compared to unrelated previews. This effect was expected to be similar for the two groups in the horizontal reading direction, but larger in the top-down direction for the Taiwanese than for the mainland Chinese group. In addition, for eye movement measures, we expected a preview effect, with fixations after identical previews being shorter than after unrelated previews. Specifically, we expected that the size of the preview benefit would depend on, both, participant group and reading direction in a three-way interaction: The preview effect was expected to be similar for the two groups in the left-right reading direction, but larger for the Taiwanese than the mainland Chinese group in the top-down direction because of the presumably larger downward perceptual span of the Taiwanese group.

All methods and proposed analyses for the experiment were pre-registered at https://osf.io/34u92/.

## 2. Methods

### 2.1. Participants

Thirty native Chinese (Mandarin) speakers, originally from mainland China (16 females; mean age = 20.5 years, SD = 2.56), and another 30 native Chinese (Mandarin) speakers, originally from Taiwan (16 females; mean age = 22 years, SD = 2.87), participated in the combined EEG/eye-tracking experiment. All participants were college students studying in Hong Kong. At the time they were recruited, participants had resided in Hong Kong for 2 years on average; the two groups did not differ in the time they had lived in Hong Kong (Mainlanders: *M* = 1.94 years, *SD* = 2.11, range: 0.17–8 years; Taiwanese: *M* = 2.07, *SD* =1.60, range: 0.08–6 years). Importantly, before coming to Hong Kong members of both groups had continuously lived in their respective home regions.

A self-report questionnaire was administered to evaluate the participants’ reading and writing experiences in horizontal and vertical directions before and after coming to Hong Kong. Participants were asked about their exposure to vertically and horizontally aligned texts in 10 different types of media: magazines, books, comics, newspapers, textbooks, contents in smart devices, road signs, billboards, slogans/leaflets and advertisements. As expected, Mainlanders reported more experience in reading horizontal texts than Taiwanese (including textbooks, slogans, leaflets, road signs; all *t*s > |2.92|, all *p*s < 0.05 for the different categories of text), whereas Taiwanese reported more experience in reading vertical texts (including textbooks, all *t*s > |3.48|, all *p*s < 0.001). In addition, as Taiwanese are usually more familiar with vertically aligned texts, 9 participants in the Taiwanese group reported convert texts in smart devices into the vertical direction through apps, whereas 7 Mainlanders reported to convert texts from vertical into horizontal direction. Taiwanese estimated to be exposed to vertically aligned text at an earlier age than Mainlanders (5.2 vs. 10.85 years old, *t*_(58)_ = 7.50, *p* < 0.001), whereas Mainlanders were exposed to horizontally aligned text earlier than Taiwanese (3.5 vs 4.5 years old, t _(58)_ = 3.73, *p* < 0.001). In addition to reading, Taiwanese participants reported having more experience in writing vertically compared to Mainlanders (both before and after moving to Hong Kong, *t*s > |2.02|, *p*s < 0.05), whereas the reverse pattern was found with regard to horizontal writing habits (*t*s > |3.57|, *p*s < 0.001). After moving to Hong Kong, Taiwanese participants still had more experience in reading vertical texts compared to Mainlanders (*t*_(58)_ = 3.17, *p* = 0.002, but not for textbooks, leaflets, road signs and slogans), whereas Mainlanders had more experience in reading horizontal texts than Taiwanese (including textbooks and road signs, *t*s > |3.15|, *p*s < 0.003, but not for slogans, leaflets, *t*s < |1.65|, *p*s > 0.11). The total time spent on reading for both groups were similar before and after coming to Hong Kong (including books, comics, magazines, newspapers and contents on smart devices). Hence, as intended, Mainlanders show dominant exposure to horizontal texts, whereas horizontal and vertical reading directions appear to be more balanced in Taiwanese participants.

All participants were right-handed, without dyslexia or ADHD, and showed normal or corrected-to-normal vision (as assessed before the experiment with the Freiburg Visual Acuity and Contrast Test; Bach, 1996). Written informed consent was obtained prior to the experiment. All participants were reimbursed with 50 Hong Kong dollars (about 7 USD) per hour. The study was approved by the Joint Chinese University of Hong Kong-New Territories East Cluster Clinical Research Ethics Committee.

### 2.2. Materials

Two-character words were selected that occur in both traditional and simplified Chinese with the same meaning in Taiwan and mainland China; hence, the visual forms of these words are identical in both regions. Words that are region-specific or representing names were excluded. Only medium- or high-frequency words in both regions were selected (mainland Chinese, WF-MC: *M* = 2.26, *SD* = 0.47, range: 1.51–3.57, retrieved from SUBTLEX-CH corpus; Cai and Brysbaert, 2010; Taiwanese Chinese, WF-TC: *M* = 1.55, *SD* = 0.63, range: 0–3.21^1^, retrieved from Sinica Corpus, Chen et al., 1996).

For the parafoveal preview manipulation at the target word position, 72 critical words were selected. These words were presented twice as post-boundary target words (once after an identical and once after an unrelated preview) and twice as parafoveal previews (once as identical preview and once as unrelated preview). To counterbalance the two reading directions and the assignment of items to a particular direction, we created two sets of words by matching the number of strokes and word frequencies in mainland Chinese and Taiwanese.

### 2.3. Construction of word lists

Target words and their previews were embedded within lists of other nouns (“fillers”). Each list consisted of five words. Specifically, to create the 5-word lists, we selected 576 filler words (72 lists × 4 words × 2 sets), which were also presented twice during the experiment. Filler words were matched with target words regarding word frequency (according to the Sinica database and the SUBTLEX-CH database) and the number of strokes per character in the first and second position. The pre-target words were of medium to high frequency (WF-MC > 2.27, WF-TC > = 0.47) and of low to medium visually complexity (stroke number < 21).

In total, we therefore created 288 (144 × 2 directions) lists consisting of one critical word and four filler words each. Words in a list were phonologically and semantically unrelated and did not orthographically overlap (no homophones, shared semantic or phonetic radicals, see below for details). The target words were placed either at list positions two, three or four; accordingly, the pre-target words were placed at list positions one, two or three. In order to avoid visual overlap between the preview and the post-boundary target word at the critical list position, the target word was always presented in a different font compared to the parafoveal preview word. This implies that the fonts of previews and pre-target words were the same. If the preceding words were presented in a Kaiti font, the following words were presented in PMingliu font, and vice versa. Two filler words following each other were presented either in the same font (50%) or in the other font (50%), precluding the usefulness of font type as a cue for the upcoming target words.

#### 2.3.1. Preview-target pairs

As a basis for constructing the critical target nouns and their respective identical or unrelated parafoveal previews we took a set of 72 pairs of Chinese two-character nouns for each reading direction (e.g., 巨星–巨星 and 巴掌–巴掌), yielding the basis for identical previews. For these identical noun pairs, 72 unrelated noun pairs were created by exchanging the preview word with a word of similar number of strokes and frequency, yielding two new pairs without any semantic or other associations (e.g., 巨星–字典 and 巴掌–池塘). In the following, such a set of an identical word pair and its unrelated recombination is called a “preview-target unit”.

#### 2.3.2. Animal lists

As animal name target words for the reading task (animal name detection), we created an additional 30 lists (15 × 2 sets), which contained the name of an animal equiprobably at one of the five list positions (cf. Dimigen et al., 2012). The embedded animal names had a mean number of 19.53 strokes (*SD* = 6.73), a mean frequency of 1.91 (*SD* = 0.43) in simplified Chinese and 1.50 in traditional Chinese (*SD* = 0.39) respectively, and were matched with the filler words in the animal lists in terms of stroke number and frequency (*t*s < 1.16, *p*s >0.26). Except for the embedded animal names, the word lists containing an animal name were indistinguishable from the lists used in regular trials. They followed the same design principles, containing the same preview manipulations (animals only in target but not in preview positions in unrelated preview trials) in the same proportions as regular trials. During the experiment, the 15 animal lists were presented randomly among the 72 target lists of each direction. Data of the animal lists was excluded from analyses.

#### 2.3.3. Balancing

Lists were constructed with the aim of minimizing the orthographic, phonological, and semantic overlap between fillers and the embedded words of the preview-target unit. Besides, there was no character sharing similar pronunciations among a given word list. Stroke numbers and word frequencies were matched between filler and target words (see Table 1). Four additional participants (two each from Taiwan and mainland China) who did not participate in the experiment rated the semantic relatedness of each word list on a scale from 1 to 5. With a mean score of 1.49 (*SD* = 0.38) and 1.45 (*SD* = 0.34) in each set, no significant difference was found for semantic relatedness. For each direction, lists were presented randomly intermixed.

**Table 1.** Similarity Measures for Targets and Fillers in the Two Sets of Stimuli.

| Measure | target | filler | $p$ | |
| --- | --- | --- | --- | --- |
| Strokes in Set 1 | 18.31 (3.57) | 17.73 (3.11) | 0.30 | *n.s.* |
| Word Frequency (SUBTLEX-CH) in Set 1 | 2.28 (0.39) | 2.23 (0.24) | 0.43 | *n.s.* |
| Word Frequency (Sinica) in Set 1 | 0.002 (0.002) | 0.0021 (0.001) | 0.22 | *n.s.* |
| Strokes in Set 2 | 18.83 (3.01) | 18.21 (3.11) | 0.23 | *n.s.* |
| Word Frequency (SUBTLEX-CH) in Set 2 | 2.25 (0.40) | 2.28 (0.27) | 0.60 | *n.s.* |
| Word Frequency (Sinica) in Set 2 | 0.002(0.002) | 0.002 (0.002) | 0.26 | *n.s.* |

*Note.* Given are means across words. Standard deviations are provided in parentheses.

For both reading directions, the target words were matched according to the number of strokes, word frequency in mainland Chinese and Taiwanese Mandarin. To equate the two sets of stimuli, the fillers in the two stimulus sets were also matched (see Table 2). Furthermore, the materials used in the two sets were counterbalanced across participants.

**Table 2.** Similarity Measures for Fillers in the Two Sets of Stimuli.

| Measure | Set 1 | Set 2 | $p$ | |
| --- | --- | --- | --- | --- |
| Target: Strokes | 18.32 (3.57) | 18.83 (3.01) | 0.35 | *n.s.* |
| Target: Word Frequency (SUBTLEX-CH) | 2.28 (0.39) | 2.25 (0.40) | 0.67 | *n.s.* |
| Target: Word Frequency (Sinica) | 0.002 (0.002) | 0.002 (0.001) | 0.49 | *n.s.* |
| Filler: Strokes | 17.73 (3.11) | 18.21 (3.11) | 0.36 | *n.s.* |
| Filler: Word Frequency (SUBTLEX-CH) | 2.24 (0.24) | 2.28 (0.27) | 0.29 | *n.s.* |
| Filler: Word Frequency (Sinica) | 0.002(0.002) | 0.002 (0.002) | 0.28 | *n.s.* |

*Note.* Given are means across words. All standard deviations are provided in parentheses.

### 2.4. Procedure

Participants were seated in a dimly-lit electrically shielded chamber at a distance of 90 cm from a monitor (24 in. BenQ ZOWIE XL2411K, resolution: 1920×1080 pixels; vertical refresh rate: 144 Hz). In two separate blocks, participants read the word lists horizontally or vertically, with a short pause within each block. The order of presentation (vertical reading first or horizontal reading first) was counterbalanced across the participants. The words in the two sets were also counterbalanced across the vertical and horizontal conditions. During the experiment, two identical monitors were used, one was oriented horizontally and one vertically for the horizontal and vertical reading directions, respectively; participants switched between these monitors between blocks. During a given reading direction block, the appropriate monitor was used, while the other monitor was moved aside. Participants were instructed to read each list and to indicate at the end whether it had contained an animal name.

The trial schemes are illustrated in Figure 1. Horizontal and vertical trials began with the presentation of a fixation cross on the left or top of the screen, respectively. After a fixation on this point was registered by the eye-tracker, the list of five words appeared on the horizontal or vertical midline, respectively. Words were presented in black on a white background. Each two-character word in the list extended a visual angle of 1.8° horizontally or vertically, depending on reading direction. In addition, there was one empty character space between the words. The visual angle between the right/lower edge of the pre-target word and the left/upper edge of the target word was 5.5°, and an invisible boundary was placed at 2.5° between words.

**Figure 1.**
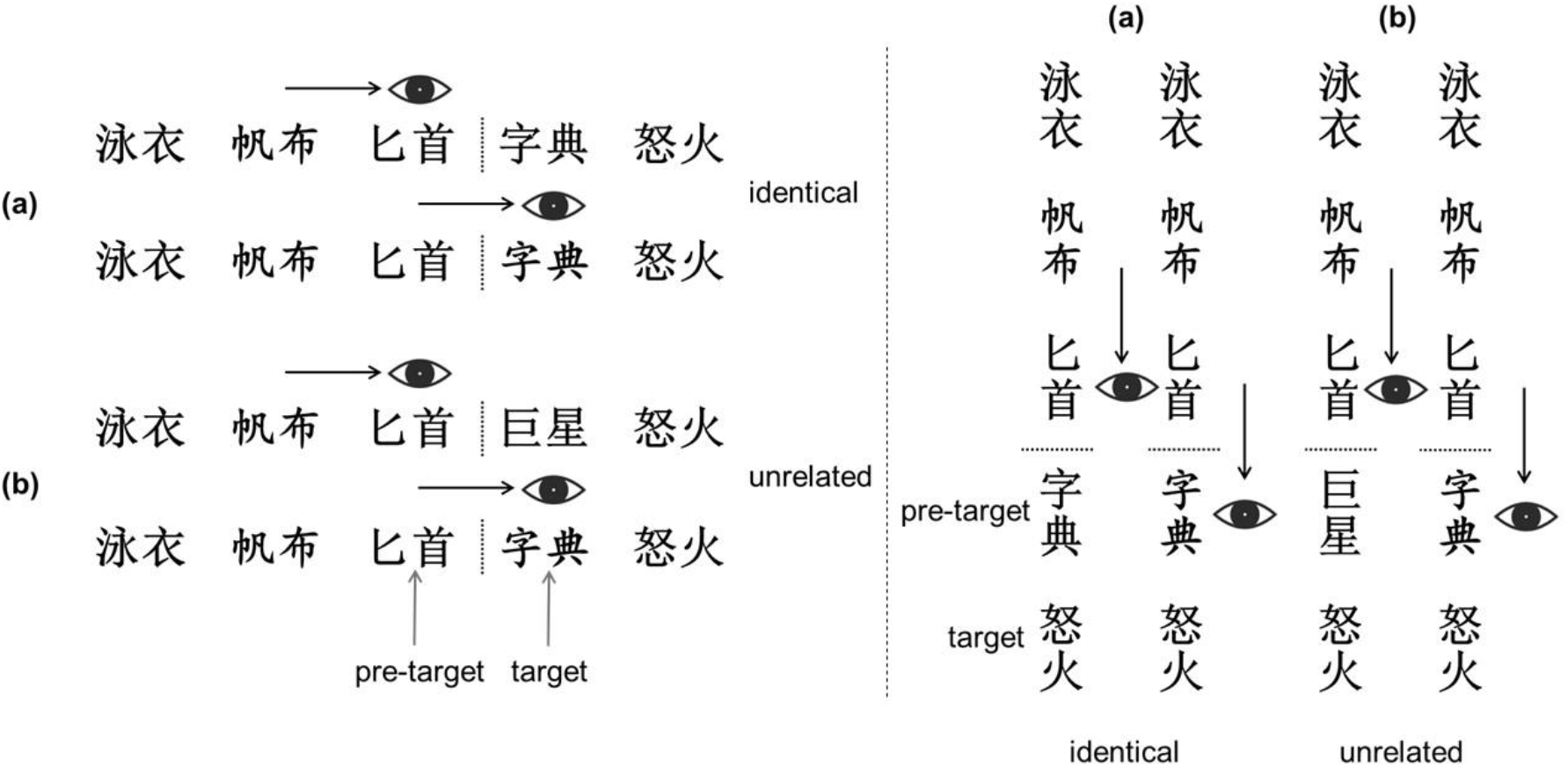
Illustration of experimental conditions. Participants read 5-word lists with the task to detect an occasional animal name in the lists. For one word in each list, the parafoveal preview was manipulated using the gaze-contingent boundary paradigm. While participants’ eyes were still looking at the pre-target word, the parafoveal preview word could be either (a) identical or (b) unrelated to the target word fixated after the saccade. Left panel: horizontal reading direction. Right panel: vertical reading direction.

**Figure 2.**
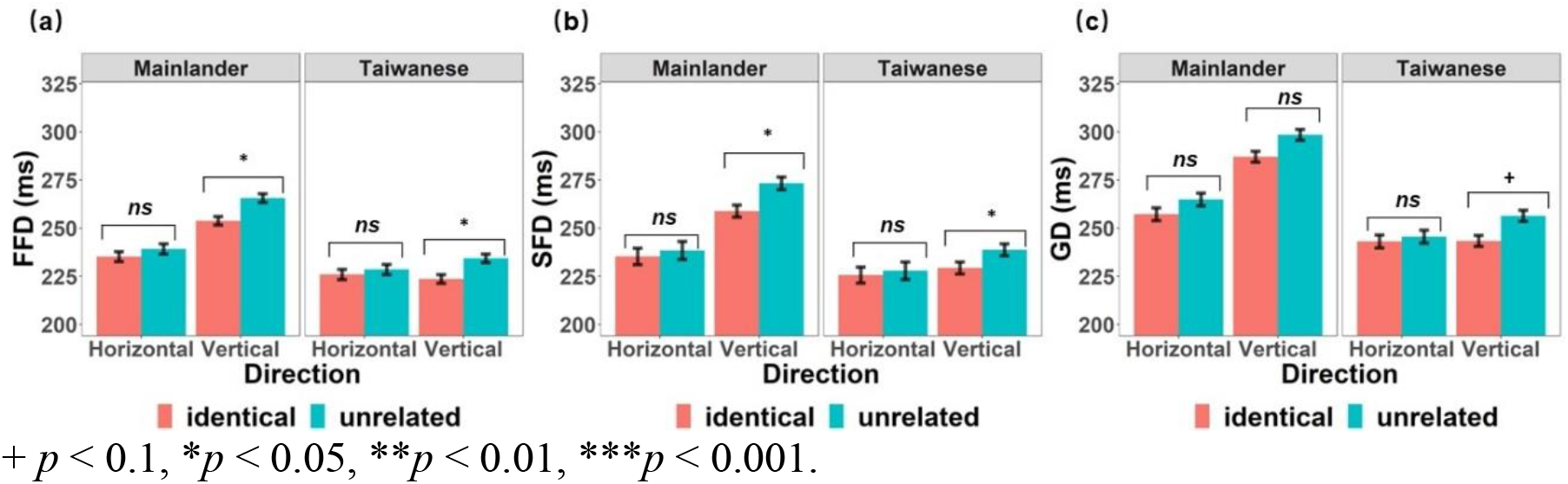
Preview effect on the target word (fixation times following an identical versus unrelated preview). Fixation durations (±1 standard error) are presented separately for FFD, SFD and GD.

As shown in Figure 1, following list onset, participants read the five words, moving their eyes freely over the text. After finishing reading, they looked at the final fixation point. After 500 ms, a blank screen appeared, and participants used two buttons to respond with left or right index fingers whether or not they had seen an animal name in the list. The assignments of yes or no responses to the left or right index finger were counterbalanced across participants. Participants read six lists for practice.

Display change awareness was assessed after the experiment. Participants were first asked whether they had noticed “anything strange about the visual display of the text” (White et al., 2005). They were asked again if they had noticed any changes if they answered “no,” after which they were informed that changes had occurred. If the answer was yes, participants were asked to (1) estimate the number of changes perceived, (2) report the identity of some of the preview words, and (3) report the list positions at which changes had occurred (see Dimigen et al., 2012).

### 2.5. EEG recording

The EEG was recorded from 64 Ag/AgCl scalp electrodes mounted in a textile cap at standard 10–5 system positions and referenced online against the CPz electrode. Two electro-oculogram (EOG) electrodes were placed on the outer canthus of each eye and one EOG electrode was placed on the infraorbital ridge of the left eye. Signals were amplified with an EEGO amplifier system (Advanced Neuro Technology, Enschede, Netherlands) at a band-pass of 0.01-70 Hz and sampled at 1000 Hz. Impedances were kept below 20 kΩ.

### 2.6. Eye movement recording

Eye movements were recorded binocularly at a sampling rate of 1000 Hz using an Eyelink 1000 plus eye tracking system (SR research) in the desktop-mounted (remote) configuration. Head position was stabilized via the chin rest of the tracker. A 9-point calibration was completed at the beginning of the experiment and before each change in reading direction. Extra calibrations were performed whenever a check failed. Calibration was accepted when the average error was < 0.5° and the maximum error < 0.99°. Furthermore, a 1-point drift correction check was performed at the beginning of each trial.

### 2.7. Co-registration of eye movements and EEG

The co-registration of eye movements and EEG was achieved by sending shared trigger pulses from the presentation PC (running Presentation, Neurobehavioral Systems Inc., Albany, CA) to the EEG and eye tracking computer on each trial through the parallel port. This allowed for accurate offline synchronization of eye movements and EEG signals via the EYE-EEG extension for EEGLAB (http://www.eyetracking-eeg.org, Dimigen et al., 2011). After synchronization, the temporal offset between the shared markers in both recordings rarely exceeded 1 ms.

### 2.8. Preprocessing of eye movement data

Three eye movement measures were used for data analysis, including first-fixation durations (FFD), single fixation durations (SFD), and gaze durations (GD). Fixations were determined by Data Viewer software (SR research). Only fixations that occurred during the first-pass reading in trials with a correct answer to the animal question were analyzed. Specifically, fixations on the area of interest were excluded, when the display change occurred too early or too late (i.e., when the display change took more than 10 ms before/after fixation onset on the target character). We also removed trials with FFD < 60 ms or > 600 ms and GD > 800 ms (total number of excluded fixations: 644). Additionally, we excluded fixations on target words in which participants blinked. In addition, we removed all trials with an incorrect manual response to the animal question. Taken together, we collected 15,724 observations for all participants. The trials left for each condition were listed in *Table.3*.

**Table 3.** Mean number of analyzed trials, standard deviations and range.

|  | Condition | *M* | *SD* | range |
| --- | --- | --- | --- | --- |
| Mainlanders | H-identical | 64.43 | 5.90 | 50–72 |
|  | H-unrelated | 66.07 | 4.68 | 57–72 |
|  | V-identical | 64.30 | 6.89 | 37–72 |
|  | V-unrelated | 64.60 | 6.28 | 41–72 |
| Taiwanese | H-identical | 62.43 | 8.27 | 30–70 |
|  | H-unrelated | 62.73 | 7.09 | 44–71 |
|  | V-identical | 61.43 | 9.06 | 39–72 |
|  | V-unrelated | 61.80 | 6.89 | 37–72 |

*Note.* H = horizontal reading direction; V = vertical reading direction.

### 2.9. EEG preprocessing

Offline, EEG data were digitally band-pass filtered, using FIR with EEGLAB 2020.0 (Delorme and Makeig, 2004) toolbox for Matlab (version 2018b), between 0.1 Hz and 30 Hz (−6 dB/octave) and re-calculated to the average reference (Lehmann and Skrandies, 1980). Independent component analysis (ICA) was used for ocular correction using procedures implemented in the EYE-EEG extension. Specifically, following the ICA decomposition, we removed all independent components that showed much more activity during saccades than during fixation periods (saccade/fixation variance ratio > 1.1) following the procedures and threshold recommendations provided in Plöchl et al. (2012) and Dimigen (2020).

After ocular correction, the EEG signal was segmented from 300 ms before to 700 ms after the first fixation onset on a word. The baseline was corrected by subtracting the 150 ms preceding the fixation onset on the target word. Epochs with amplitudes exceeding ±100 μV in any channel (except the EOG) were automatically rejected from further analyses. FRPs were then averaged within and then across participants.

After eye movement and FRP preprocessing, across all 60 participants, our screening left us with a total number of 7,348 good epochs for the target character in the vertical reading condition and 5,741 good epochs in the horizontal reading condition. Within each reading direction, there was a similar numbers of remaining epochs for the unrelated and identical previews (unrelated, *M* = 48.47, *SE* = 1.03; identical, *M* = 48.31, *SD* = 1.08). However, the number of remaining trials was significantly different for the two reading directions (main effect, *t* _(59)_ = −13.38, *p* < 0.001) because participants failed the trial-initial fixation check more often in the horizontal than in the vertical direction. The analysis of variance (ANOVA, with Bonferroni correction on post-hoc tests) on the number of remaining trials in the two groups and directions showed that neither the main effect of Group nor the interaction with trial number were significant (*F*s < 1.58, *p*s > 0.21)

### 2.10. Data analysis

#### 2.10.1. Eye movements

Eye movement data were analyzed with linear mixed-effects models (LMMs) within the *R* environment for statistical computing (R Core Team, 2015). We estimated variance components for subjects and for items (i.e., varying intercepts and slopes), using the “lmer” function of the *lme4* package (Bates et al., 2015; version 1.1.27.1) on log transformed FFDs, SFDs and GDs. The within-subject factors of *Preview* (identical vs. unrelated) and *Direction* (vertical vs. horizontal) and the between-subject factor *Group* (Taiwanese vs. Mainlander) were coded as fixed factors. Participants and items were specified as crossed random effects, with both random intercepts and random slopes (Barr et al., 2013). When we ran the models, we always began with full models that included the maximum random effects structure. But the slopes were removed if the model failed to converge (indicating over-parametrization). The *p*-values were estimated using the “lmerTest” package with the default Satterthwaites’s method for degrees of freedom and *t*-statistics (Kuznetsova et al., 2017).

#### 2.10.2. FRP data analysis

We analyzed FRPs time-locked to the first fixation onsets on the target words by using LMMs. The analysis was preregistered (https://osf.io/34u92/), including time windows and selected electrodes. As previous studies mainly selected a time window of 200–280 ms (e.g., Dimigen et al., 2012) or a time window of 180–280 ms (Buonocore et al., 2020), and Kornrumpf et al. (2016) suggesting that the preview positivity emerged earlier than 200 ms after fixation onset, we selected 180–280 ms as the time window of the preview positivity. Besides, as many studies observed an early N1 component, we selected 120–160 ms as the time window of the early N1^2^. As the early N1 effects and preview positivity have a scalp distribution over occipito-temporal regions, we selected this area as region of interest (ROI; left occipital-temporal area, LOT: PO9/PO7, and right occipital-temporal area, ROT: PO8/PO10) and also included a factor of *Hemisphere* (left vs. right). The same LMM statistics as for eye movements were applied to FRP epochs, except the factor *Hemisphere* was also included as additional predictor. Post-hoc analyses were performed to obtain contrasts, and the tests were adjusted using the multivariate *t*-distribution (mvt) in the *emmeans* package (Lenth, 2019; version 1.7.3).

## 3. Results

### 3.1. Display change awareness

In the post-experimental debriefing about the awareness of saccade-contingent preview manipulation, all participants, except one in the Taiwanese group, were aware of changes of words from the preview to the target. Eight Mainlanders were able to correctly report the positions of previews located in the vertical reading direction, and 9 could do so for the horizontal direction. Four Taiwanese correctly reported the position of previews in the vertical direction and 6 Taiwanese correctly reported it for the horizontal direction. In addition, a question about the estimated number of previews showed that Mainlanders reported more changes in the horizontal than the vertical direction (*M* = 37.1 vs. 34.7); in contrast, Taiwanese noticed more changes in the vertical than in the horizontal direction (*M* = 26.2 vs. 24.7). These findings indicate that participants were not able to recognize the previews, and the prevalence of display change awareness seems to be similar across the two participant groups, although almost all of them noticed the saccade-contingent preview manipulation.

### 3.2. Animal task performance

On average, participants detected 97.7% of the animal names contained in the lists (*d*’ = 3.92), as shown in Table 4. The *d-primes* were calculated for each participant and each direction separately. Repeated measures ANOVAs on the within-subject factor direction and the between-subject factor group showed that *d-primes* did not differ as a function of either factor (*Direction, F* _(1, 58)_ = 1.67, *p* = 0.20; *Group, F* _(1, 58)_ = 1.52, *p* = 0.22; *Direction × Group, F* _(1, 58)_ = 0.08, *p* = 0.78). These animal task results suggest that readers read the word lists attentively for comprehension in both groups, regardless of reading direction.

**Table 4.** Means (and standard deviations, in parentheses) of Sensitivity (d’) for Responses to Targets in Vertical and Horizontal Reading Directions between Mainlanders and Taiwanese Group.

|  | Mainlander | Taiwanese |
| --- | --- | --- |
| Horizontal (SD) | 3.78 (0.54) | 3.93 (0.50) |
| Vertical SD) | 3.93 (0.46) | 4.02 (0.53) |

### 3.3. Eye movements

The eye movement data showed that reading times on the target words following identical previews were shorter than after unrelated previews, confirming the classic preview effect (see Table 5). This preview effect was significant for FFD (difference of 8 ms), and GD (difference of 9 ms) but not for SFD (difference of 7 ms). The vertical reading lists required longer fixation durations than the horizontal lists in terms of FFD (difference of 13 ms), SFD (difference of 19 ms) and GD (difference of 19 ms). Taiwanese were generally slightly faster readers than Mainlanders, with shorter FFDs (difference of 20 ms), SFDs (difference of 21 ms) and GDs (difference of 29 ms). In addition, we observed significant interactions between *Group* and *Direction* for FFD, SFD and GD, with a larger FD difference between the horizontal and vertical reading directions in Mainlanders than in Taiwanese. Finally, the interaction between *Preview* and *Direction* was significant for FFD and marginally significant for GD (see Table 6 for the results of the linear mixed-effects model), such that preview effects were larger in the vertical than in the horizontal direction. However, the three-way interaction between *Preview, Direction* and *Group* was not significant for any of the three eye movement measures. Therefore, behaviour provided no evidence of group differences in preview effects in the two reading directions, although both groups tended to show larger preview effects in vertical than in horizontal direction.

**Table 5.** Means and Standard Errors of the Fixation Time Measures (in Milliseconds) in the Different Conditions

|  | Condition | FFD | SFD | Gaze |
| --- | --- | --- | --- | --- |
| Mainlanders | H-identical | 233 (18) | 235 (18) | 257 (23) |
|  | H-unrelated | 237 (19) | 238 (18) | 265 (24) |
|  | V-identical | 253 (14) | 258 (13) | 287 (20) |
|  | V-unrelated | 265 (16) | 273 (15) | 299 (21) |
| Taiwanese | H-identical | 225 (17) | 225 (17) | 243 (24) |
|  | H-unrelated | 228 (18) | 228 (18) | 245 (22) |
|  | V-identical | 223 (15) | 229 (15) | 243 (21) |
|  | V-unrelated | 234 (16) | 239 (16) | 256 (20) |

*Note.* FFD = first fixation duration; SFD = single fixation duration; GD = gaze duration; H = horizontal reading direction; V = vertical reading direction.

**Table 6.** Fixed Effect Estimates from the Linear Mixed-Effects Models on the Eye Movement Data

| Factor | First fixation duration |  |  |  | Single fixation duration |  |  |  | Gaze duration |  |  |  |
| --- | --- | --- | --- | --- | --- | --- | --- | --- | --- | --- | --- | --- |
|  | *b* | *SE* | *t* | Sign. | *b* | *SE* | *t* | Sign. | *b* | *SE* | *t* | Sign. |
| (Intercept) | 5.400 | 0.013 | 341.279 | <0.001*** | 5.420 | 0.018 | 299.992 | <0.001*** | 5.470 | 0.017 | 316.627 | <0.001*** |
| Direction | 0.074 | 0.020 | 3.659 | 0.001** | 0.086 | 0.023 | 3.735 | <0.001*** | 0.096 | 0.007 | 12.745 | <0.001*** |
| Group | -0.088 | 0.007 | -13.138 | <0.001*** | -0.088 | 0.011 | -8.410 | <0.001*** | -0.116 | 0.008 | -15.393 | <0.001*** |
| Preview | 0.025 | 0.01 | 2.643 | 0.009** | 0.023 | 0.014 | 1.637 | 0.11 | 0.027 | 0.010 | 2.562 | 0.01* |
| Group × Direction | -0.094 | 0.001 | -6.991 | <0.001*** | -0.104 | 0.021 | -4.955 | <0.001*** | -0.108 | 0.015 | -7.196 | <0.001*** |
| Direction × Preview | 0.031 | 0.001 | 2.319 | 0.02* | 0.033 | 0.020 | 1.6 | 0.11 | 0.027 | 0.015 | 1.819 | 0.069+ |
| Group × Preview | 0.002 | 0.001 | 0.161 | 0.87 | -0.002 | 0.020 | -0.099 | 0.92 | 0.001 | 0.015 | 0.079 | 0.93 |
| Preview × Group × Direction | 0.01 | 0.03 | 0.363 | 0.72 | -0.010 | 0.041 | -0.254 | 0.80 | 0.027 | 0.030 | 0.904 | 0.37 |

2 + $p < .1$ . \* $p < .05$ . \*\* $p < .01$ . \*\*\* $p < .001$ .

### 3.4. EEG results

Figure 3 shows the grand-average FRPs, time-locked to the first fixation on target words. The visual inspection on the visual forms at OT electrodes showed the biphasic muscle spike potential around time zero (Keren et al., 2010), followed by a P1-N1 complex. This complex consisted of the P1 component peaking around 100 ms after fixation onset, and an early N1 component peaking around 170 ms. The early N1 peak showed larger amplitudes for identical previews than unrelated ones. After the early N1 peak, the FRP amplitude during the falling flank of the N1 (the late N1) was substantially larger for identical previews than unrelated ones.

**Figure 3.**
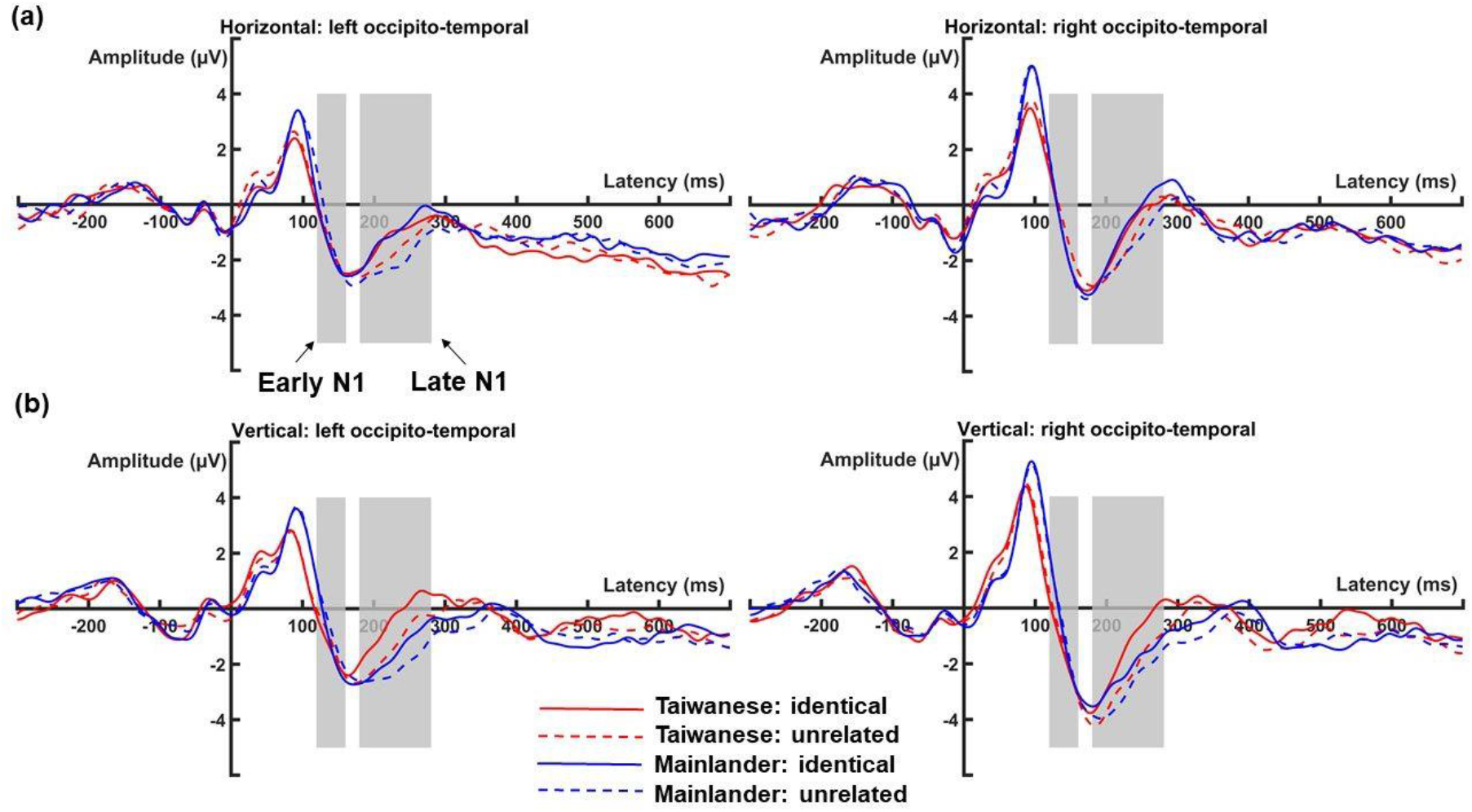
Fixation-related potential (FRP) waveforms for the horizontal (a) and vertical reading direction (b) at the left and right hemisphere occipito-temporal regions of interest (LOT and ROT). Gray regions mark the a priori-defined time windows used for the analyses of early N1 and late N1.

#### 3.4.1. LMMs results (preregistered analyses)

##### Early N1

The early N1 component (120–160 ms), tended to be larger (more negative) after identical than unrelated previews, although the effect was only a trend (*Preview*, *b* = 0.53, *SE* = 0.29, *t* = 1.81, *p* = 0.07, see Figure 4). In addition, this preview effect was not different between the two groups (*Preview × Group, b* = −0.40, *SE* = 0.38, *t* = −1.06, *p* = 0.29) or the two reading directions (*Preview* × *Direction, b* = −0.34, *SE* = 0.35, *t* = −0.38, *p* = 0.71). The three-way interaction between *Preview, Group* and *Direction* was also not significant (*b* = 0.29, *SE* = 0.50, *t* = 0.59, *p* = 0.56). In addition, the early N1 amplitudes were more negative in the left hemisphere than in the right hemisphere (*Hemisphere*, *b* = 0.79, *SE* = 0.26, *t* = 2.99, *p* = 0.003). All other main effects and interactions were not significant.

**Figure 4.**
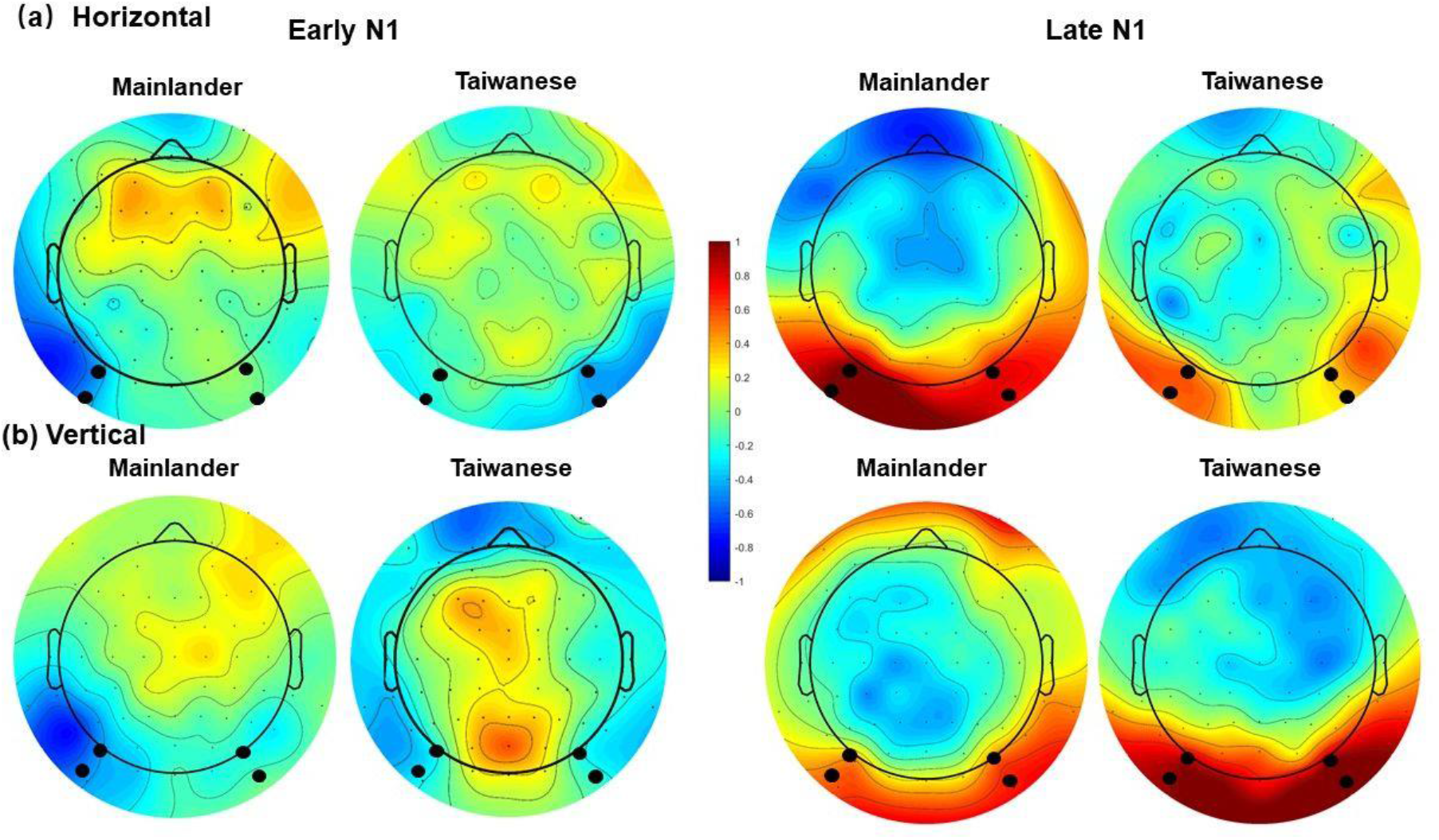
The topographies of the preview effect (identical minus unrelated) for the horizontal (a) and vertical (b) reading direction for Mainlander and Taiwanese. Topographies are shown for the early N1 (left side) and late N1 (right side). Black dots highlight the electrodes used to define the regions of interest (LOT and ROT).

**Figure 5.**
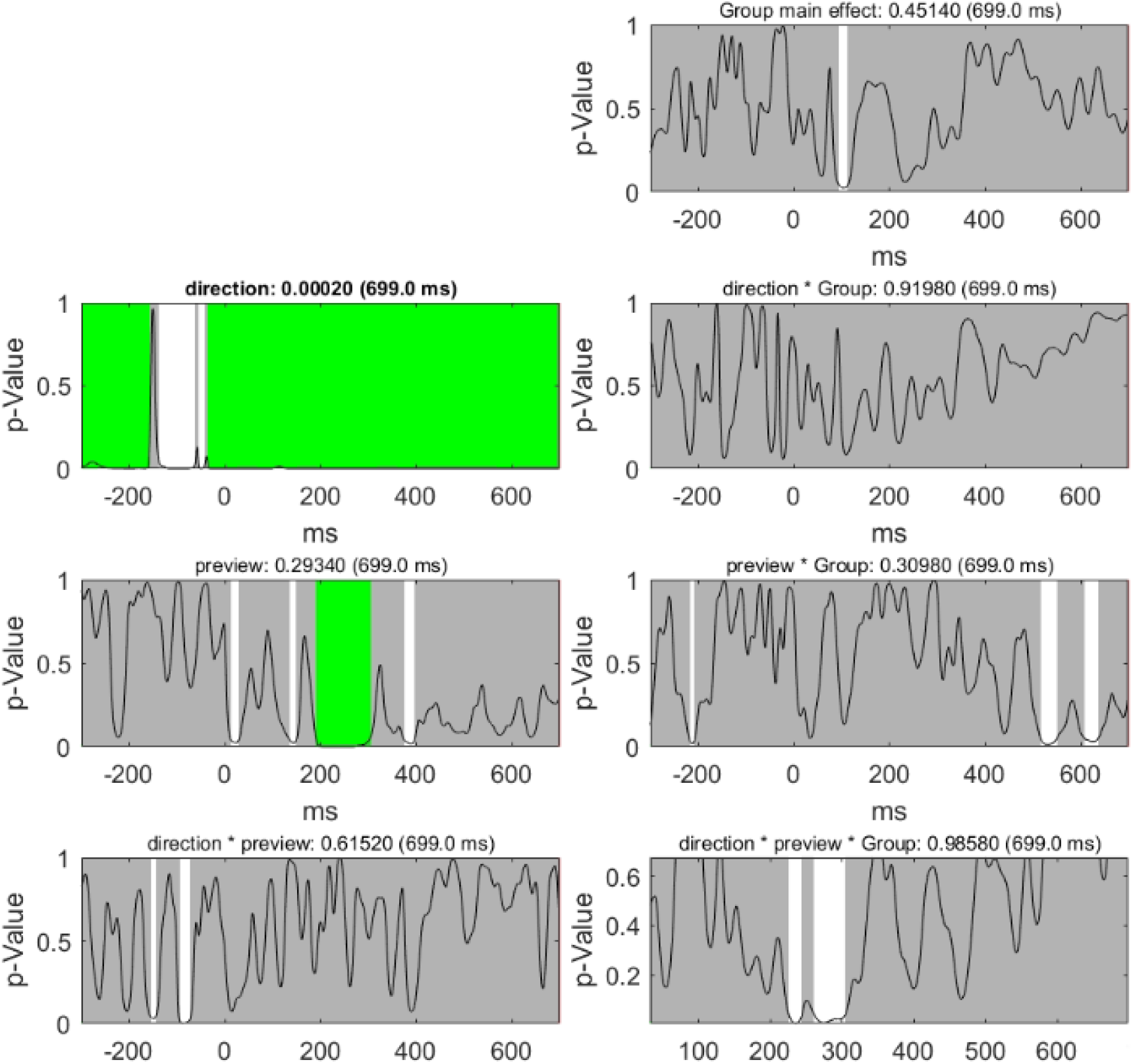
Results of the exploratory sample-by-sample TANOVA for the different factors and contrasts. Each plot visualizes the *p*-values (y-axis) for the comparison between the mean FRP maps of each factor level or interaction for every time point after fixation onset (milliseconds on the x-axis). Gray areas mark non-significant time points, whereas the white areas mark periods of significant differences between the factor levels. We corrected for multiple comparisons using global duration statistics, and the duration thresholds were then applied to the TANOVA plots. These periods longer than the estimated duration threshold are marked in green (i.e., effects corrected for multiple comparisons). For the main effect of *Direction*, the duration threshold was identified as 82 ms. For the main effect of *Preview*, the duration threshold was identified as 45 ms.

##### Late N1

In the late N1 time window (180–280 ms), we found a reduced (i.e., more positive) FRP amplitude following identical as compared to unrelated previews (*Preview*, *b* = −0.99, *SE* = 0.30, *t* = −3.28, *p* = 0.001, see Figure 4), replicating the previously reported “preview positivity” effect. Importantly, this preview effect differed between the Taiwanese and Mainlander groups as a function of reading direction, as indicated by a significant three-way interaction between *Group, Preview* and *Direction* (*b* = −0.93, *SE* = 0.46, *t* = −2.00, *p* = 0.046). Post-hoc analyses revealed that Taiwanese showed a significant preview effect only in the vertical direction (*b* = 0.97, *SE* = 0.23, *z* = 4.20, *p* < 0.001) whereas Mainlanders showed a significant preview effect only in the horizontal direction (*b* = 0.86, *SE* = 0.25, *z* = 3.47, *p* = 0.003). Also, in the horizontal direction, FRP amplitudes were similar for both groups, whereas in the vertical direction the amplitudes were more negative for Mainlanders than for Taiwanese (*Direction* × *Group, b* = 1.29, *SE* = 0.63, *t* = 2.04, *p* = 0.04). In addition, the FRP amplitudes were more negative in the left hemisphere than in the right hemisphere in the horizontal direction, but the pattern was slightly opposite in the vertical direction (*Direction* × *Hemisphere, b* = −0.86, *SE* = 0.32, *t* = −2.67, *p* = 0.008). All other effects were not significant (*t*s < |1.53| and *p*s > 0.12).

#### 3.4.2. Exploratory analysis

To further investigate whether there were effects in the FRPs beyond the time windows and electrodes that we had selected a-priori, we used a sample-by-sample Topographic Analyses of Variance (TANOVA) that includes all electrodes in the FRP map. The Ragu software (Koenig et al., 2011) was used on non-normalized (raw) topographic maps to test for effects of the two within-subject factors (*Direction* and *Preview*) and the between-subject factor (*Group*). The TANOVA was corrected for multiple comparisons through Global Duration Statistics (Koenig et al., 2011). If this global test was significant (*p* < 0.05 at 5000 randomization runs), *t*-maps (across participants and against zero) of the covariance maps were computed and displayed. As we were mainly interested in preview effects corresponding to the early N1 and late N1 effects, we focused on the first 400 ms after fixation onset.

As shown in Figure 3, the main effect of *Direction* was significant across the entire time window after fixation onset^3^, and the preview effect was significant in the time window of 191–303 ms. In addition, the main effect of *Group* was significant between 93 to 115 ms. The three-way interaction for *Preview*, *Direction* and *Group* reached significance during the time window of 225–305 ms (separated by a short interval of 15 samples where *p*-values were smaller than 0.1), although the two time windows did not survive correction for multiple comparisons. The other interactions were not significant within 400 ms after fixation onset. Overall, the TANOVA results were consistent with the ROI analysis of the late N1, especially the time windows identified by TANOVA for preview effect and the three-way interaction, had a large overlap with the ROI analysis. However, for the early N1 component, the time window identified for preview effects (135–147 ms) did not survive the multiple comparison correction, and no significant time window was identified for the three-way interaction (*Preview × Direction × Group*).

## 4. Discussion

The present study investigated the influence of experience in reading in different directions, namely, horizontal and vertical, on neural correlates of visual word recognition.

We recruited participants from Taiwan and mainland China, all speaking Mandarin, and tested them on the same reading materials, presented in both horizontal and vertical directions. We used a boundary paradigm together with the co-registration of EEG and eye movements, allowing readers to move their eyes freely as in natural reading. In the boundary paradigm, either identical or unrelated previews were presented. It was expected that the preview effects would differ as a function of different reading directions between the two groups of readers, especially in the vertical direction, due to the much more pervasive experience of Taiwan residents with vertical script.

Results replicated several common observations in behavior. First, we found typical preview effects in eye movements. In addition, both groups performed very accurately in the animal detection task, suggesting that reading performance was good and not significantly different between groups. More importantly, not only did we find the typical preview positivity, its presence in the vertical reading direction depended on the prior experience with reading in that direction. The main findings of the preview effects in both eye movements and FRP, and how vertical reading experiences modulate preview effects are discussed below.

### 4.1. Preview effects in FRP

The analysis of fixation-related potentials (FRPs) showed a reduced occipito-temporal negativity after identical preview in the time window from 180–280 ms, which we called late N1, which in previous papers has also been called “preview positivity” (Dimigen et al., 2012) due to the more positive-going amplitudes after valid previews. In addition, we also obtained this preview effect in TANOVAs, where the significant time windows overlapped with the ones we had pre-registered, suggesting that the effect is robust and can be detected across the entire map. The preview positivity is usually considered to reflect the preview-based facilitation of early stages of visual word recognition at visual and/or orthographic levels (Niefind and Dimigen, 2016). The effect’s time course and scalp distribution (largest over left occipito-temporal regions) fit to previous late N1 or N250 findings (e.g., Bentin et al., 1999; Maurer et al., 2005), which have been linked to orthographic processing.

However, contrary to our hypothesis, we did not obtain an electrophysiological preview effect for the early part of the N1 component (preregistered as the interval between 120–160 ms), although there was a trend that identical words were more negative than unrelated words in FRP amplitudes. Although the early N1 preview effect was rarely reported in the FRP literature (e.g., Degno et al., 2019a; Dimigen et al., 2012), an N1 effect with larger negativity for primed vs. unprimed words is frequently seen in masked priming studies (Chauncey et al., 2008; Huang et al., 2022). Therefore, this early N1-like preview effect, similar to the N1 effects in the masked priming paradigm, may reflect the initial stage of sublexical orthographic processing in visual word recognition (Grainger and Holcomb, 2010). However, as the preview effect in the early N1 component was only marginally significant, the initial stage of sublexical orthographic processing may be not as robust as in the later stage (i.e., the late N1 component) of visual word recognition.

### 4.2. Does vertical reading experience modulate preview effect in FRP?

The key finding of the current study is the three-way interaction between *Preview, Group* and *Direction* for the late N1 component, which fits the reading experience hypothesis as predominant reading experience of Taiwan participants in the vertical direction (confirmed by the questionnaire) modulated the preview effects in the two directions. To be more specific, we found that Taiwanese showed larger preview effects in the vertical direction compared to the horizontal direction. This three-way interaction was not only found in the ROI analysis, but also with TANOVAs, suggesting the effect can be detected across the entire map. The three-way interaction indicates that in the vertical direction, readers with a rich vertical reading experience (i.e., Taiwanese) are better at making use of parafoveal information compared to readers that are less accustomed to vertical reading (i.e., Mainlanders).

It is also noteworthy that the timing of the preview positivity (at the level of the late N1) fits to previous studies, which found perceptual expertise effects in the N170 components in bird-experts looking at birds (Tanaka and Curran, 2001), and in car-experts looking at cars (Gauthier et al., 2003), and also for individuals with expertise in print words (Maurer et al., 2005), and for humans looking at faces (for review see Rossion and Jacques, 2011). The current findings therefore suggest that long-term cultural experiences may shape the way readers process visual words, for example, by putting readers who have more exposure to vertical reading directions at an advantage in processing vertically aligned texts compared to readers who are less accustomed to this reading direction.

An alternative explanation for the three-way interaction could also be that the two groups have different levels of word form familiarity. Since the two-character words were arranged differently in the vertical and horizontal directions, it renders their visual word forms also different in the two directions. As Taiwanese readers are familiar with both vertically and horizontally aligned texts and are also more familiar with words with vertical arrangements compared to Mainlanders, the resulting (un)familiarity with the visual forms during vertical reading may also contribute to the three-way interaction.

Contrary to our expectations, we did not observe any significant interactions between the early N1 preview effect with *Group* or *Directions*; this could be due to the nonsignificant early N1 preview effect. The results showed that at this early stage of visual word recognition, readers from the two groups differing in vertical reading experience may have nevertheless processed the words similarly in both directions. The results suggest that at the early stage of visual word reading, neither the visual inputs from different directions nor the readers’ reading experience in vertical direction have any major influence on the word recognition process.

### 4.3. Lateralization for reading direction?

We also obtained an interaction between *Direction* and *Hemisphere* in the late N1 component such that the vertical direction showed a right-hemisphere bias but the horizontal direction showed a slight tendency towards left-lateralization in both groups. Previous literature has found that the reading-related N170 is left-lateralized (Bentin et al., 1999; Maurer et al., 2005; Tarkiainen et al., 1999), with larger amplitudes over the left hemisphere for words than for low-level visual stimuli. This left-lateralized N170 topography elicited by visual words stands in contrast to N170 responses for other forms of perceptual expertise related to faces or objects of expertise, which are typically bilateral or right-lateralized (Rossion et al., 2003; Tanaka and Curran, 2001). Previous hypotheses suggested that left-lateralization of the N170 was due to the involvement of phonological processing during learning to read (phonological mapping hypothesis; Maurer and McCandliss, 2007) or due to a larger degree of high spatial frequencies in visual words (spatial frequency hypothesis, Mercure et al., 2008). However, neither of these two hypotheses can explain the current findings with left-lateralization only for the horizontal direction, as phonological influences and spatial frequencies were the same for the two reading directions. This finding is potentially very interesting, and suggest another or additional mechanism that may explain left-lateralized processing of visual words.

The left-lateralization for visual words is considered to be part of the left hemisphere language network, and this left-hemispheric dominance for language has been found to be not only associated with alphabetic languages (e.g., Brem et al., 2006; Cohen et al., 2000), but also logographic languages (i.e., Japanese, Maurer et al., 2008; Chinese, Tan et al., 2001; Xue et al., 2019). However, evidence for left-lateralization for printed script is usually derived from writing systems in which the text runs from left to right, while for scripts with a right-left orientation, the lateralization is sometimes right-biased but further neuroimaging evidence is lacking (e.g., Hebrew, Yiddish, for a review, see Obler, 1989; but in Orbach, 1952). Therefore, the right-laterization for words during vertical reading in the current study may suggest left hemisphere activation for visual-orthographic information may be related to left-to-right reading, and therefore could be related to eye movements and attention allocation. As in left-right reading, the parafoveal information is located in the right visual field, which may further influence visuospatial attention and oculomotor behavior. However, further investigation is needed to test these hypotheses.

### 4.4. No modulation of behavioural preview effects by vertical reading expertise

Consistent with the FRP analysis, the eye movement data also showed typical preview effects, as the preview effects were significant in both FFD and GD, consistent with previous reports (Buonocore et al., 2020; Degno et al., 2019b, 2019a; Dimigen et al., 2012). Preview effects in fixation times were small (e.g., 8 ms in FFD) compared to those in previous studies (e.g., FFD: 20 ms in Dimigen et al., 2012; 38 ms in Dimigen and Ehinger, 2021; 41 ms in Degno et al., 2019a; 35 ms in Yan et al., 2009; 14 ms Yang et al., 2009; see Tsang and Chen, 2012 for a review). The small size of the preview effect in the current study may be at least partly due to the use of word lists rather than sentences as materials. As we used word lists as materials, it was not possible to predict upcoming words based on sentence context, which likely facilitates preview effects during sentence reading.

We also found that both Taiwanese and Mainlanders fixated longer during vertical reading than horizontal reading. This finding is consistent with previous reports, as Yan et al. (2018) found that Taiwanese showed longer fixation durations in vertical than horizontal reading. A similar finding was obtained for readers without expertise in vertical reading (Laarni et al., 2004), indicating that the vertical reading experiences may not modulate fixation durations on left-right and top-down reading directions.

In addition, some biological factors may have also an influence on the reading direction effects. For example, the spatial density of photoreceptor cell along the horizontal direction is generally higher than along the vertical direction (Curcio et al., 1990). Similarly, Najemnik and Geisler (2008) found that target visibility drops faster vertically than horizontally. Furthermore, evidence on the neurological control of horizontal and vertical eye movements has shown that vertical saccades are slower than horizontal saccades, and the downward saccades are the slowest (Terry Bahill and Stark, 1975). Therefore, the biological basis and the neural control of the visual system may influence reading on the vertical and horizontal directions of texts.

The eye movement data did not show the key three-way interaction between *Preview, Direction* and *Group*, which we observed in FRPs. The absence of the three-way interaction in eye movements is likely a consequence of the numerically small preview effects in the groups for both directions. Although the three-way interaction was not significant, the two-way interaction between *Group* and *Direction* was, as Taiwanese had shorter fixation durations in vertical directions than Mainlanders, indicating that the vertical reading expertise modulates fixation durations. Alternatively, the absence of the three-way interaction in eye movements but its presence in FRPs may suggest that FRPs are more sensitive to the preview effect than eye movements. Fixation-related potentials show high temporal resolution and can reflect on-line processes, whereas fixation durations only capture the summed duration of all cognitive processing occurring during word identification. Therefore, it is possible that preview effects reached largest positivity before the current fixation is completed.

In addition, we obtained a main effect of *Group*, with Mainlanders showing overall longer fixation durations than Taiwanese, although comprehension performance (as indicated by the performance in the animal task) was not significantly different. This overall group effect cannot be explained by the faster vertical reading in the Taiwanese group, as further test on each reading direction showed longer fixation durations for Mainlanders than Taiwanese even in the horizontal direction. A possible reason may be that readers who use simplified Chinese system (i.e., Mainlanders) processed the text in a less holistic way than traditional Chinese readers (i.e., Taiwanese) when perceiving characters (Liu et al., 2016), therefore Mainlanders may be more sensitive to internal constituent components of characters and may need more time for recognition. Such long-term influences of reading and writing experience with the two writing systems cannot be ignored, although the materials we selected have the same visual forms in both simplified and traditional Chinese.

### 4.5. Limitations

Our study also has several potential limitations. First, we noted that the number of accepted trials for the FRP analysis differed between the reading directions, with more remaining data for the vertical direction (due to a smaller number of failed fixation-checks at the beginning of the trial). However, we believe that the fewer trials in the horizontal direction cannot explain the critical three-way interaction, as the two groups did not differ from each other with regard to the trial number in a given reading direction. Especially for the FRP analysis, no two-way interaction of *Group* and *Direction* was found, and Mainlanders showed a preview effect only in the horizontal but not in the vertical direction, suggesting no weakening of the preview effect in the horizontal direction.

Second, the differences of writing systems may have an impact on readers’ character perception. Consistent evidence has shown greater visual discrimination skills in readers of simplified rather than traditional Chinese (Mcbride-Chang et al., 2005; Peng et al., 2010; Yang and Wang, 2018) and more analytic character processing (Liu et al., 2018). Therefore, readers of the two writing systems may process the same characters in different ways, which may further influence the neural correlates of character processing. However, this cannot explain the three-way interaction, as such an effect should be independent of the reading direction.

Third, all participants in the current study were living in Hong Kong at the time of the study, where the horizontal reading direction is mainly used. Our Taiwanese participants had therefore presumably less exposure to vertical text as compared to readers who were residing in Taiwan more recently. The observed preview effects in the vertical direction for the Taiwanese group in the current study may therefore be a conservative estimate of the influence of the experience with a reading direction, and it may also partially explain the absence of the three-way interaction between *Group, Direction* and *Preview* in the eye-movement data. Further studies may address this issue by recruiting participants that were not recently exposed to a different proportion of reading directions.

## 5. Conclusions

The present study provides the first evidence that readers with vertical reading expertise show a larger preview positivity in their fixation-related brain activity compared to readers that are less accustomed to vertical text. This modulation of preview effects by the reader’s cultural experience indicates that long-term reading experience in the vertical direction shapes the way how readers process written words. Future studies may consider the potential influences of reading experiences on different stages of visual word processing.

## Acknowledgements

This work was supported by the Germany/Hong Kong Joint Research Scheme of the Research Grants Council of Hong Kong (G-CUHK409/18) and the German Academic Exchange Service (DAAD, Project 57447990).

## Declaration of Competing Interest

The authors declare that they have no known competing financial interests or personal relationships that could have appeared to influence the work reported in this paper.

## Footnotes

1 Please note that the word frequencies for the same words in Sinica Corpus are usually lower than in SUBTLEX-CH corpus.

2 Note that previous papers did not always differentiate between the early N1 component and the preview positivity in their analysis, but sometimes referred to the preview positivity as the “late parts” or “falling flank” of the N1 component (Kornrumpf et al., 2016). Our current analysis distinguished between these two intervals. The two components were distinguished by the direction of preview effects.

3 We noted that for the main effect of *Direction*, all *p*-values were smaller than 0.05 after fixation onset, while in the ROI analysis, the main effect of Direction was not significant. Possibly, the selection of electrodes in the ROI cannot explain the direction differences as the selection of the ROI was based on preview effects. Further t-maps on the effect of *Direction* showed that the largest activation in the corresponding time windows of the early and late N1 component was located in the eye electrodes and central-posterior sites, thus, the occipito-temporal electrodes may not reflect the *Direction* effect.

## Notes

https://osf.io/34u92/

### Competing Interest Statement

The authors have declared no competing interest.

